# Lineage-specific accumulation of endogenous dsDNA viral elements across eukaryotes

**DOI:** 10.1101/2025.04.19.649669

**Authors:** Hongda Zhao, Lingjie Meng, Ruixuan Zhang, Morgan Gaïa, Hiroyuki Ogata

**Author notes:** Corresponding author: Hiroyuki Ogata.

## Abstract

Endogenous viral elements record past virus–host interactions, yet those derived from Baltimore class I double-stranded DNA viruses remain poorly explored in eukaryotes. Here, we identified 781,111 viral regions in 7,103 eukaryotic genome assemblies, revealing their broad but uneven distribution across both eukaryotic host lineages and viral taxa. Viral regions showed pronounced accumulation in several invertebrates, where they accounted for up to 16% of a single genome assembly. Taxonomic analyses of viral regions revealed 72 previously unrecognized endogenization-based associations between viral major taxa and eukaryotic lineages, including those involving invertebrates such as bivalves, starfish, sponges, flatworms, and ribbon worms, as well as diverse algae. In addition, we uncovered cases of iridoviral endogenization in ray-finned fishes. Overall, our study provides a baseline view of the landscape of the endogenous dsDNA virosphere across both eukaryotic and viral lineages.

## Introduction

Endogenous viral elements (EVEs) are widespread across diverse eukaryotic genomes and provide genomic footprints of past virus–host encounters^1,2^. Endogenous elements derived from reverse-transcribing viruses, including those from Baltimore class (BC) VI, are known to account for a considerable proportion of certain eukaryotic genomes^3^. For example, endogenous retroviruses alone constitute ∼8% of the human genome^4^. These viral elements can cause disease, provide essential host functions through ‘co-option’ or contribute antiviral functions to the host^5–7^.

In contrast to reverse-transcribing viruses, endogenous elements derived from BC I (dsDNA) viruses remain underexplored^2,8^. A few eukaryotic BC I viruses, such as human herpesvirus 6 and the mavirus virophage, are known to integrate their genomes into host chromosomes during their replication cycles^9,10^. Additional BC I viral lineages have recently been shown to persist within host cells as integrated or episomal forms, with the potential for reactivation and virion production^10–13^. Domesticated dsDNA viral elements can enhance host survival, as exemplified in parasitoid wasps, where *Bracoviriform* particles derived from EVEs are delivered into parasitized lepidopteran larvae to suppress immunity and support wasp development^14,15^. Recent studies have begun to uncover signatures of specific BC I virus lineages in algal genomes^16–18^ and horizontal transfers of viral genes to eukaryotes^19–21^.

Eukaryotic BC I viruses are polyphyletic and highly diverse. They range from small viruses, such as papillomavirids^22^ (including papillomaviruses) and viruses from the polinton-like supergroup^23^, to giant representatives like megaviricetes^24^. Of note, the known eukaryotic dsDNA virosphere is rapidly expanding owing to culture-independent metagenomics^25^, which has led to discoveries of novel dsDNA viral groups, including polinton-like viruses^26^ (PLVs, also classified as *Aquintoviricetes*), *Mirusviricota*^27^, *Mriyaviricetes*^28^, and the proposed *Egovirales* order^29^. Eukaryotic genome data are also accumulating at an impressive pace through international initiatives such as the Earth BioGenome Project^30^. Therefore, it is timely to re-evaluate dsDNA viral elements in diverse eukaryotes in light of these expanding datasets to better understand the endo-virosphere.

## Results and Discussion

### Baltimore class I viral regions are widespread but unevenly distributed across eukaryotic lineages

We conducted a systematic analysis to detect dsDNA VRs across eukaryotic genomic assemblies. In total, 37,254 assemblies from GenBank, covering diverse lineages as of Feb. 2024 were analyzed (**Fig. 1a, Supplementary Table S1**). Our study incorporated 2,126 reference viral genomes from 11 BC I viral taxa known or predicted to infect eukaryotes (**Table 1, Supplementary Table S2**), which are referred to here as viral major taxa.

**Fig. 1.**
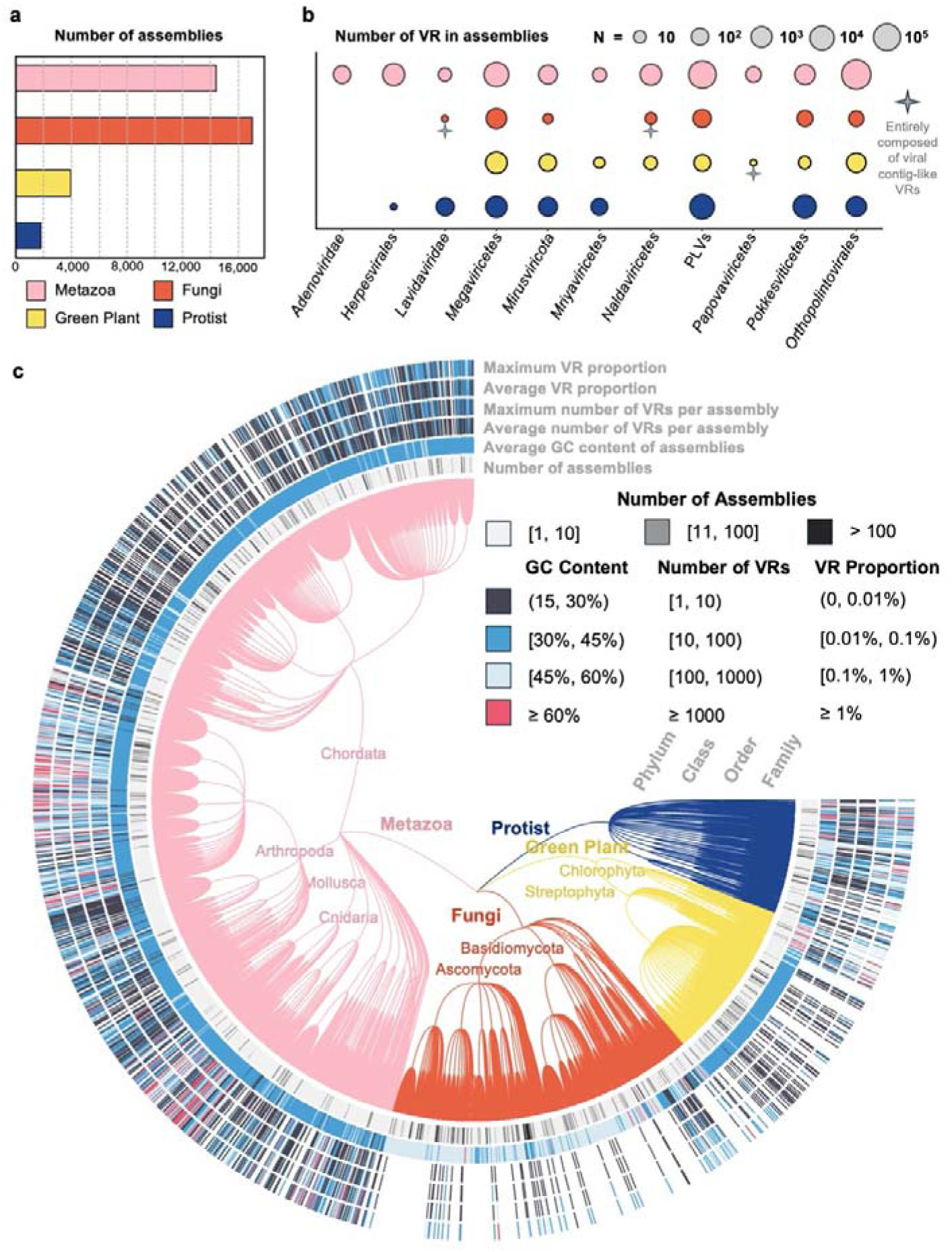
Viral regions identified in eukaryotic genome assemblies. (a) The number of eukaryotic genome assemblies used in this study. Colors indicate different eukaryotic groups. (b) The number of VRs identified using HMM profiles derived from reference viruses. Colors match those in (a). The radius of each circle represents the logarithmically transformed count of VRs. (c) VR content across the tree of eukaryotes. The cladogram is based on taxonomy, and each leaf represents a eukaryotic family. The attributes shown in the outer ring are explained in the diagram in the upper-right corner.

**Table 1.**
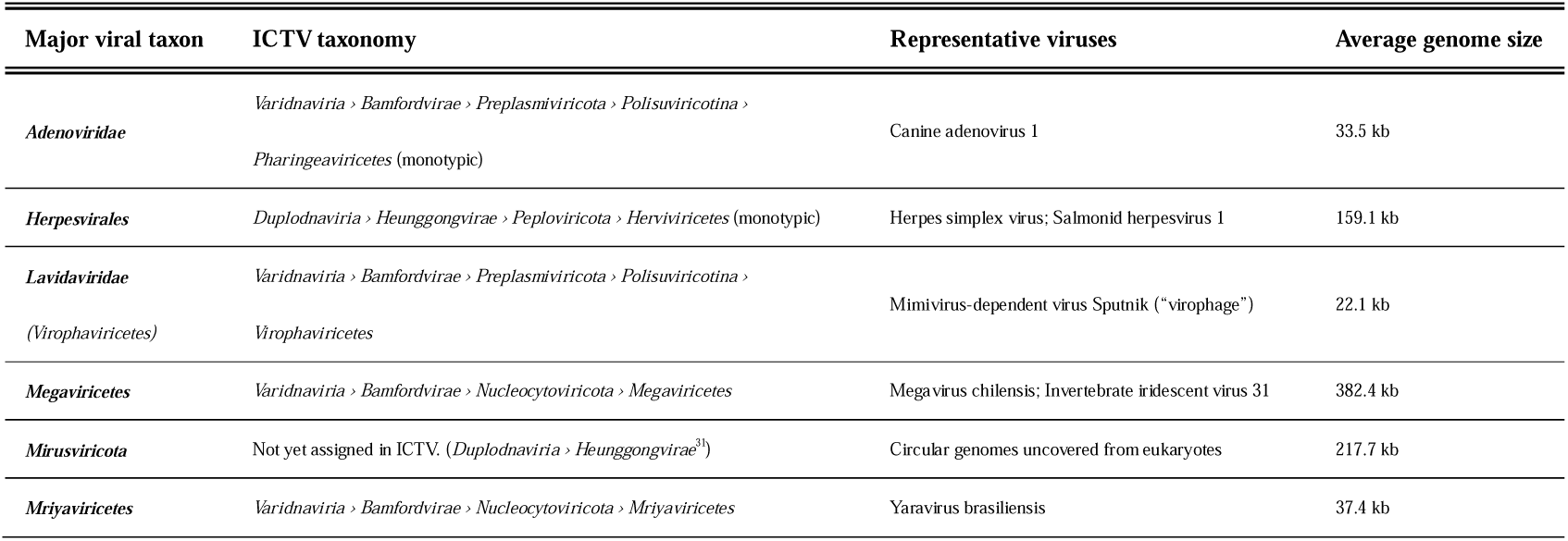
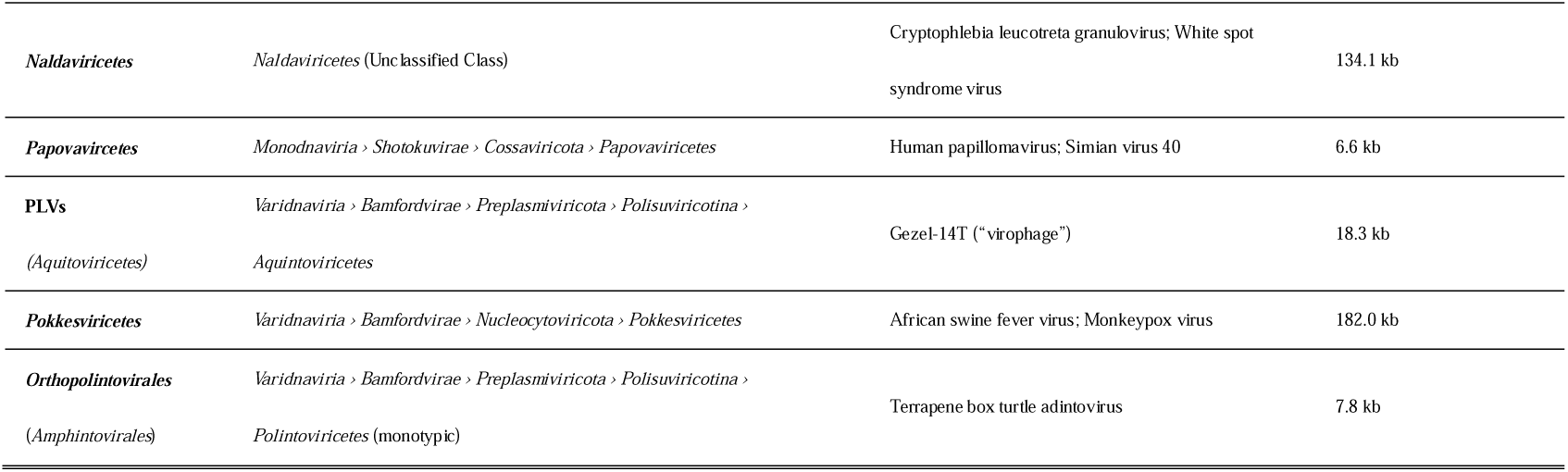
Eleven Baltimore class I viral major taxa included in this study.

This analysis identified a total of 781,111 VRs enriched in clustered viral open reading frames (ORFs) across 7,103 genome assemblies (19% of all assemblies). These included 6,094 metazoan assemblies (42.25% of all metazoan assemblies) and 633 protist assemblies (34.39%). We preliminarily assigned each identified VR to a viral major taxon based on the HMM profiles used to detect it. The majority of VRs were taxonomically associated with *Orthopolintovirales* and PLVs (**Fig. 1b, Supplementary Table S3**). Even though no BC I viruses have been isolated from fungi or higher plants, 168 Streptophyta assemblies (4.52%) and 153 fungal assemblies (0.89%) were found to contain dsDNA VRs, including those preliminarily assigned to *Megaviricetes* and *Orthopolintovirales*. These results are consistent with previous observations of imiterviral (*Megaviricetes*) signals in a bryophyte, a lycophyte^31,32^, and zoosporic fungi^33^.

The distribution of VRs across different host lineages was uneven, with metazoans accounting for the largest proportion (757,529 VRs, 96.98%), followed by protists (20,186 VRs, 2.58%), green plants (2,442 VRs, 0.31%), and fungi (954 VRs, 0.12%) (**Fig. 1b**). When the calculation was restricted to VR-containing assemblies, the mean of family-level average VR numbers per assembly was 101.84, 19.23, 1.72, and 0.28 for metazoans, protists, green plants, and fungi, respectively. VRs were identified in 2,951 genera, but approximately 80% of VRs were derived from 300 genera (**Supplementary Fig. S1a**), and 4.3% originated from a single genus, *Unio* (bivalve). We examined the relationship between the abundance of VRs and the completeness of genome assemblies (**Supplementary Fig. S1b**). Neither the VR content (i.e., the proportion of VRs in individual assemblies) nor the number of VRs was correlated with assembly completeness, indicating that potential viral contamination in low-quality assemblies had no major influence on the overall detection of VRs.

To distinguish potentially integrated viral sequences from unintegrated viral sequences, we classified VRs into ‘integration-like’ and ‘viral contig-like’ categories based on the VR and contig lengths as well as the cellular OG content. The majority (n = 766,069; 98%) of VRs were classified into the integration-like category, whereas only a small subset of VRs (n = 15,042; 2%) were classified as viral contig-like and lacked signatures of integration.

### General features of identified viral regions

Several characteristics of VRs were examined to assess their quality as *bona fide* virus-derived regions. Regions derived from viral genomes are expected to be enriched in coding sequences compared with eukaryotic genomic regions. The ORF density of a VR or a eukaryotic assembly was calculated as the total length of ORFs divided by the length of the VR or the assembly, respectively. The analyzed VRs exhibited higher coding potential than their host genomes (**Supplementary Fig. S1d)**, with most showing 20–40 percentage points higher, suggesting an origin distinct from that of the surrounding genomic regions.

Tetranucleotide frequency (TNF) differences between VRs and their host genomic regions were also examined (see Methods for details). The proportion of VRs showing TNF compositions distinct from the host genomes was highly dependent on their assigned viral major taxa (**Supplementary Fig. S1f)**. VRs preliminarily assigned to *Herpesvirales*, *Papovaviricetes*, and *Lavidaviridae* tended to exhibit TNF compositions distinct from their host assemblies, whereas most VRs related to *Orthopolintovirales*, *Pokkesviricetes*, and *Mriyaviricetes* displayed compositions more similar to those of their hosts.

The cellular homologs within the identified VRs was further assessed to further evaluate their viral origin (**Supplementary Fig. S1c**). Nearly three-quarters of VRs contained less than 20% of their ORFs with best hits to cellular reference sequences, indicating a general enrichment of viral genes in the VRs.

ORFs in VRs were shorter than those in the reference viral genomes. The expected length range of a viral ORF was defined based on the length distribution of ORFs in the same OG from reference viral genomes of the same viral major taxon. In most VRs, nearly half of their ORFs fell outside this range (**Supplementary Fig. S1e)**, suggesting that pseudogenization through degradation is a common phenomenon in VRs.

We observed that some VRs exhibited chimeric signatures from multiple viral taxa (**Supplementary Data S1**). For example, the previously identified *Nucleocytoviricota*-like giant EVE in a fungal chromosome (CP080821.1)^34^ corresponded to four VRs. One of the VRs contained *Pokkesviricetes*-specific homologs and *Megaviricetes*-specific homologs. These observations may reflect either unknown viral clades with chimeric features or the accumulation of multiple independent viral insertions within the same locus.

### Identified viral regions constitute a substantial proportion of genome content in diverse eukaryotes

VR content showed substantial variation across host lineages (**Fig. 1c**). The VR content of metazoans ranged from trace levels to over 10% (**Fig. 2**). Previously unrecognized high levels of VR content were observed in invertebrate species belonging to different taxa, such as Bivalvia (*Unio delphinus*, 16.20%; *Sinohyriopsis cumingii*, 13.38%; *Unio pictorum*, 11.71%), Insecta (*Neoaliturus tenellus*, 14.68%), and Anthozoa (*Ricordea florida*, 12.90%). Most of their VRs were preliminarily assigned to PLVs and *Orthopolintovirales* (**Supplementary Table S1**). The three bivalve species belong to the same freshwater mussel family Unionidae^35^. VRs accounted for 0.004% (75 VRs from 59 assemblies) of the human genome assemblies on average, and those in vertebrate genomes accounted for 0.014% on average, with the highest proportion (0.725%) observed in the assembly of a fish, *Gasterosteus aculeatus*, which contained 14 VRs preliminarily assigned to PLVs on its chromosome Y.

**Fig. 2.**
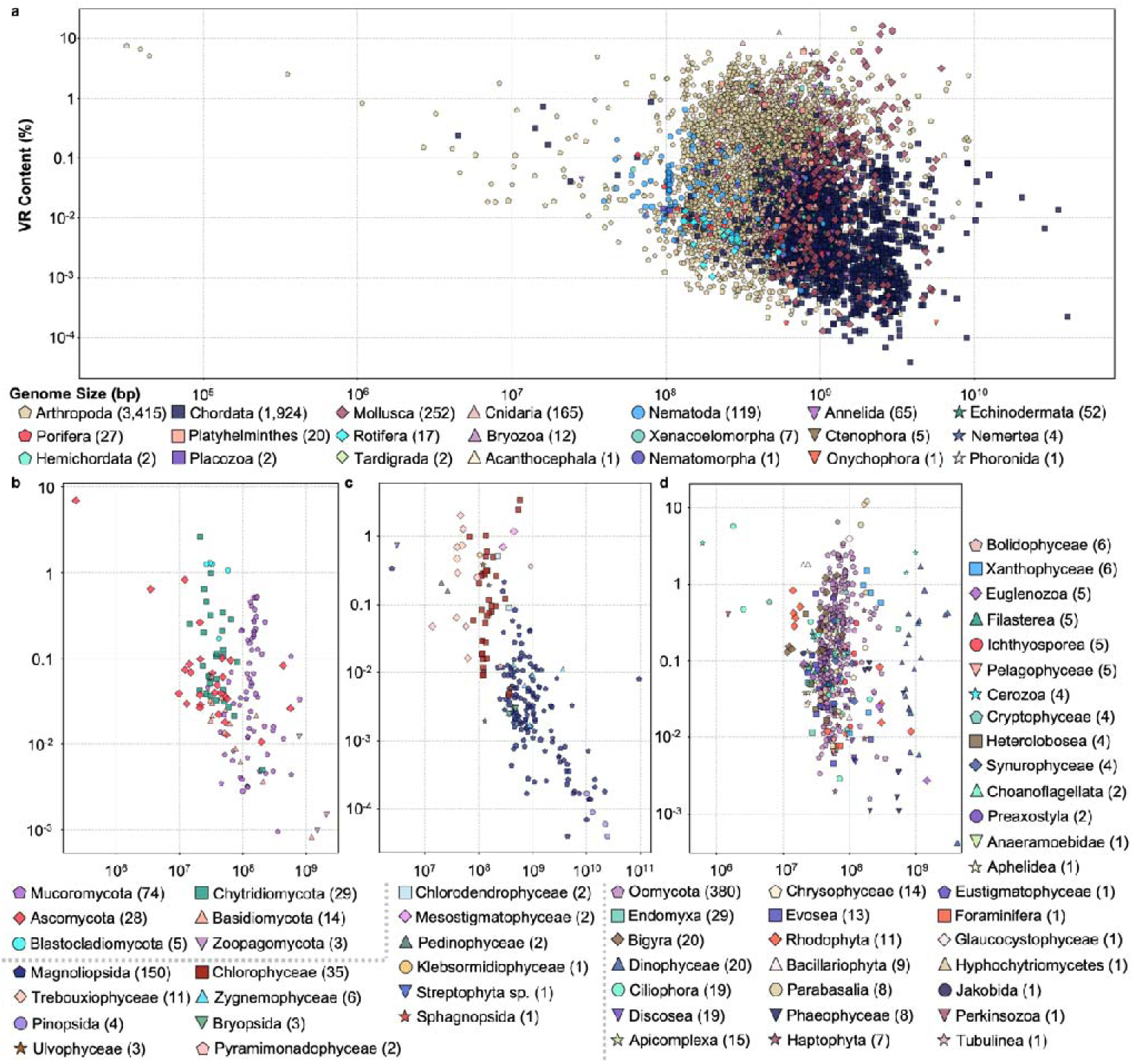
The proportion of identified VRs within different eukaryotic groups. (a–d) metazoans, fungi, green plants and protists. Each dot represents a genomic assembly, with its color and shape corresponding to its taxonomic group. The number shown in each label of eukaryotic group indicates the total number of assemblies (dots) included in the figure. Groups with fewer assemblies are plotted in the upper layer of the plot to enhance their visibility. The horizontal axis represents the assembly size, while the vertical axis represents the percentage of the genome occupied by VRs, both of which are log10-scaled.

In fungi and streptophytes, the VR content was generally low and seldom exceeded 1% (**Fig. 2**). Nevertheless, a few assemblies showed relatively high levels of VR content. Among fungi, the highest VR content was observed in a lichen assembly of *Cladonia squamosa* (6.93% driven by a single PLV VR). Among streptophytes, the unicellular biflagellate alga *Mesostigma viride* (1.19%, mainly contributed by 222 *Megaviricetes* VRs).

Protists with VR content higher than 1% belonged to Parabasalia, Oomycota, Ciliophora, Cercozoa, Bacillariophyta, Glaucocystophyceae, Dinophyceae, Xanthophyceae, and Bigyra. *Trichomonas vaginalis*, a flagellated protozoan pathogen (phylum Parabasalia), showed the highest VR content (12.13%) among protists. The *T. vaginalis* genome was enriched in VRs preliminarily assigned to *Pokkesviricetes* (1,514 VRs, 97%, closely related to *Egovirales*^29^) or, to a lesser extent, *Megaviricetes* (43 VRs, 3%).

Polintons/Mavericks are large self-replicating DNA transposons that typically encode a protein-primed family B DNA polymerase and a retrovirus-like integrase^36,37^, and many also carry virus-like morphogenesis genes such as capsid proteins and DNA-packaging ATPases^38,39^. Thus, Polinton/Maverick- and PLV-related elements blur the boundary between DNA transposons and viruses and have the potential to proliferate within host genomes. Because many VRs in high-VR-content assemblies were assigned to *Orthopolintovirales* and PLVs, high VR content could in principle reflect recent transposon-like amplification of a limited number of elements. We therefore assessed the proportion of nearly identical VRs within each assembly, defined as VR pairs sharing ≥90% identity and ≥90% coverage. Among the 435 assemblies with VR content greater than 1%, only 10 assemblies had more than 30% nearly identical VRs, with the maximum observed in the insect genome of *Sphinx pinastri* (50%) (**Supplementary Table S4**). These assemblies were enriched in VRs preliminarily assigned to *Orthopolintovirales* and PLVs, suggesting that recent transposon-like proliferation explain high VR content in a limited number of cases. In contrast, the proportion of nearly identical VRs was below 20% in 400 assemblies, including the three bivalve species with VR content greater than 10%. These results suggest that although some VRs may have recently proliferated within host genomes, high VR content in most assemblies cannot be explained solely by recent amplification of nearly identical elements.

### Occurrence patterns of Baltimore class I viral regions across eukaryotic lineages

To identify VRs with concordant viral taxonomic signals and thus infer more reliable associations between viral and eukaryotic lineages, we applied more stringent criteria (see Methods for details). VRs that passed these filters and showed taxonomic signals concordant with their initial assignments were defined as taxonomically high-confidence viral regions (HCVRs). This filtering yielded 49,584 HCVRs that showed taxonomic signals concordant with their initial assignments in metazoans, 361 in fungi, 344 in green plants, and 240 in protists. We reconstructed phylogenetic trees comprising HCVR-derived and reference viral sequences using representative marker genes conserved among reference genomes for each viral major taxon (**Supplementary Fig. S2**). These phylogenetic trees generally showed clear grouping of HCVR-derived sequences with reference viruses, supporting the taxonomic assignments of the HCVRs.

Using 50,529 HCVRs, we observed 135 occurrence-based pairs between viral major taxa and eukaryotic classes, spanning six eukaryotic supergroups (**Fig. 3**)^40^. Among these pairs, 45 overlapped with EVEs previously reported in the literature (**Supplementary Table S5**). We further compared these HCVR-based associations with virus–host relationships known from isolation studies. Of the 43 currently known associations between viral major taxa and eukaryotic classes, HCVRs captured 28 associations. After excluding associations already supported by literature or isolated viruses, 72 HCVR-based associations represented previously unrecognized virus–eukaryote links, to the best of our knowledge.

**Fig. 3.**
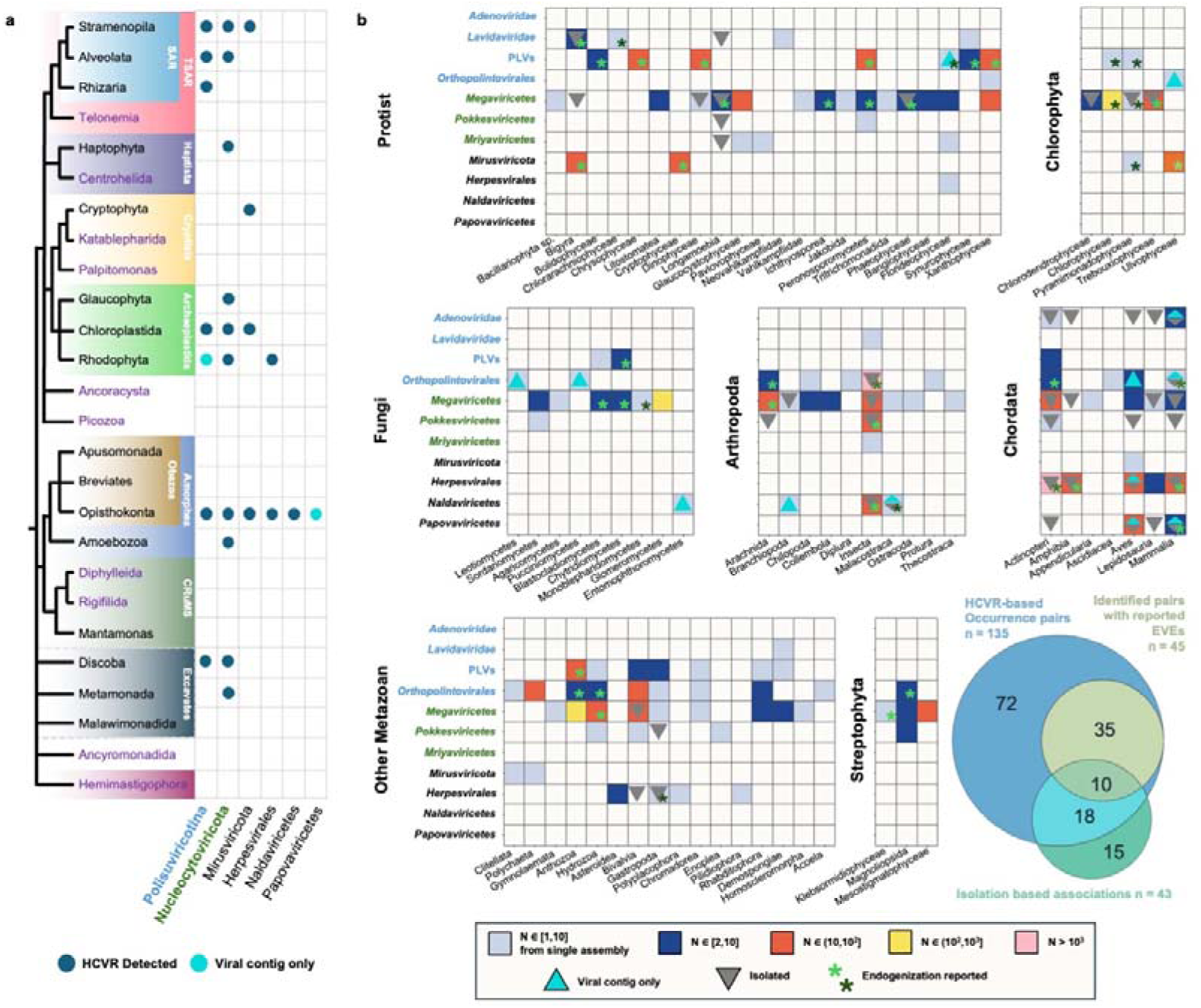
Distribution of HCVRs in eukaryotes. (a) The distribution of HCVRs across the eukaryotic tree of life. *Polisuviricotina* includes *Adenoviridae*, *Lavidaviridae*, PLVs, and *Orthopolintovirales*. *Nucleocytoviricota* includes *Megaviricetes*, *Pokkesviricetes,* and *Mriyaviricetes*. Taxa shown in purple represent lineages for which no genome assemblies are currently available in GenBank. **(b) The distribution of HCVRs in distinct eukaryotic groups.** In each heatmap, the horizontal axis represents the class-level eukaryotic taxa (or lower-level eukaryotic taxa when class-level assignments were unavailable), the vertical axis represents the viral taxa, and the color gradient indicates the number of HCVRs in each unit. A light blue triangle marks units where all HCVRs are viral contig-like, while a gray triangle indicates the presence of viruses previously isolated from these eukaryotes. The asterisk indicates that evidence of endogenization has been reported in the literature.

### Megaviricetes-related endogenization in ray-finned fish genomes

To further examine whether the occurrence-based virus–host associations reflect *bona fide* endogenization, we manually inspected the genomic loci of HCVRs representing the 135 pairs. We identified several compelling cases in which HCVRs were embedded within eukaryotic genomes and surrounded by host-derived genomic regions.

We identified 69 HCVRs assigned to *Megaviricetes* from 13 Actinopterygii assemblies, representing 12 species across nine orders (**Fig. 4a**). In fish genomes, BC I virus-derived EVEs have mainly been reported to be associated with alloherpesvirus-related lineages (**Supplementary Fig. S2**)^41^, including endogenous herpesvirus-like elements such as Teratorn^42^. By contrast, although iridovirids are pathogens of fish and related signals have been detected in fish viromes or metatranscriptomic datasets^43,44^, their endogenization in fish genomes has not been clearly recognized. The present study found that several *Megaviricetes*-related HCVRs were embedded within chromosomal regions of *Syngnathus scovelli, Cyclopterus lumpus*, and *Pelmatolapia mariae* (**Fig. 4b**).

**Fig. 4.**
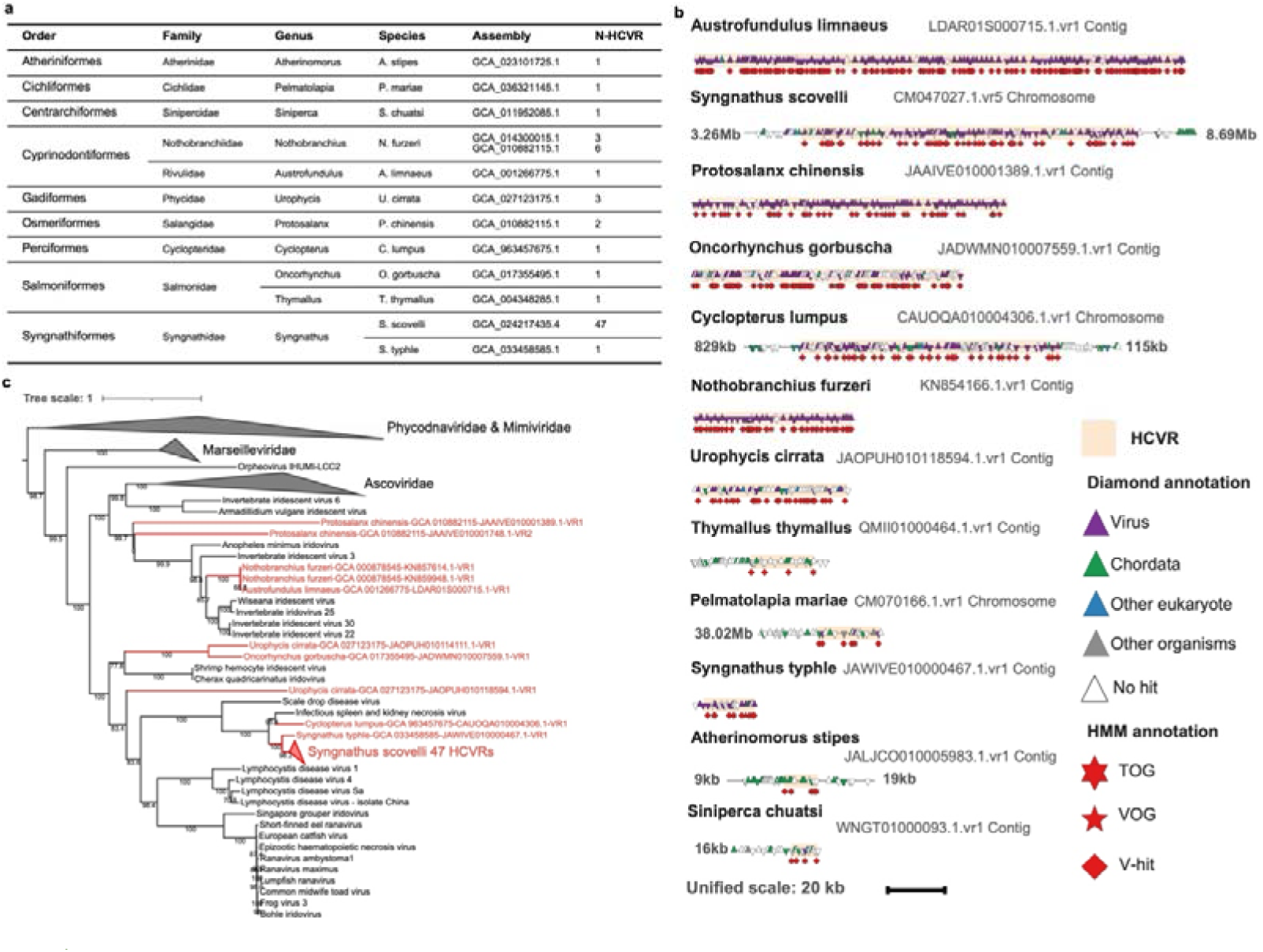
Megaviricetes-related HCVRs in actinopterygian assemblies. **(a) Taxonomic information and numbers of *Megaviricetes* HCVRs in actinopterygian assemblies. (b) Genomic maps of HCVRs in each Actinopterygii assembly.** For each map, 20 kb of flanking sequence on both sides of each HCVR is shown; when the available flanking sequence was longer than 20 kb, the unshown length is indicated as text. HCVRs are highlighted with a light-red background. Each triangle represents an ORF; upward and downward triangles indicate ORFs encoded on the positive and negative strands, respectively. Triangle colors correspond to the taxonomic affiliations of the best hits. HMM-based annotations are shown below each map. **(c) Reconstructed phylogenetic tree of HCVRs and reference megaviricetes.** The tree is midpoint rooted. Ultrafast bootstrap values are shown on branches. The best-fit model was WAG+F+I+G4.

Using five conserved *Megaviricetes*-related OGs, including ribonuclease, helicase, primase, dynein-like β chain, and myristoylated membrane protein, we reconstructed phylogenetic relationships of 57 HCVRs from nine species across seven orders (**Fig. 4c**). These HCVRs were phylogenetically placed within *Iridoviridae*, with some sequences closely related to Infectious spleen and kidney necrosis virus. Notably, although the HCVR from *Syngnathus typhle* occupied most of its contig and therefore did not provide strong evidence for endogenization on its own, it was closely related to HCVRs from *Syngnathus scovelli* that were embedded in different chromosomes.

Together, the genomic evidence for integration, the broad distribution of HCVRs across diverse ray-finned fish, and their consistent phylogenetic placement within *Iridoviridae* suggest that iridovirus-related sequences may occasionally become endogenized in vertebrate genomes. These findings provide unequivocal evidence of *Megaviricetes*-related endogenization in vertebrates, a phenomenon that has not been clearly recognized previously.

### Embedded high-confidence viral regions reveal additional putative virus–host associations

Bivalve genomes showed the highest VR content among the surveyed eukaryotic lineages (**Fig. 2**). Among the 107,631 VRs identified in bivalve genomes, 58.5% were preliminarily assigned to *Orthopolintovirales* and 40.8% to PLVs. Because *Polisuviricotina*-related VRs are widely distributed across eukaryotic assemblies (**Supplementary Data S1**) and their encoded ORFs are frequently annotated as eukaryotic proteins, distinguishing these elements from host genes and resolving signals among closely related viral groups can be challenging. We identified 71 *Orthopolintovirales* HCVRs from 10 assemblies representing nine species (**Supplementary Fig. S3a**), including clear cases embedded within host genomic regions (**Fig. 5a**). Virus-derived ORFs with best hit sequences annotated as eukaryotes were common among these HCVRs (**Supplementary Fig. S3b**); however, phylogenetic analysis supported the taxonomic assignment of these HCVRs (**Supplementary Fig. S3c**). These high-confidence cases support the interpretation that *Polisuviricotina* VRs in bivalve genomes represent genuine virus-derived components.

**Fig. 5.**
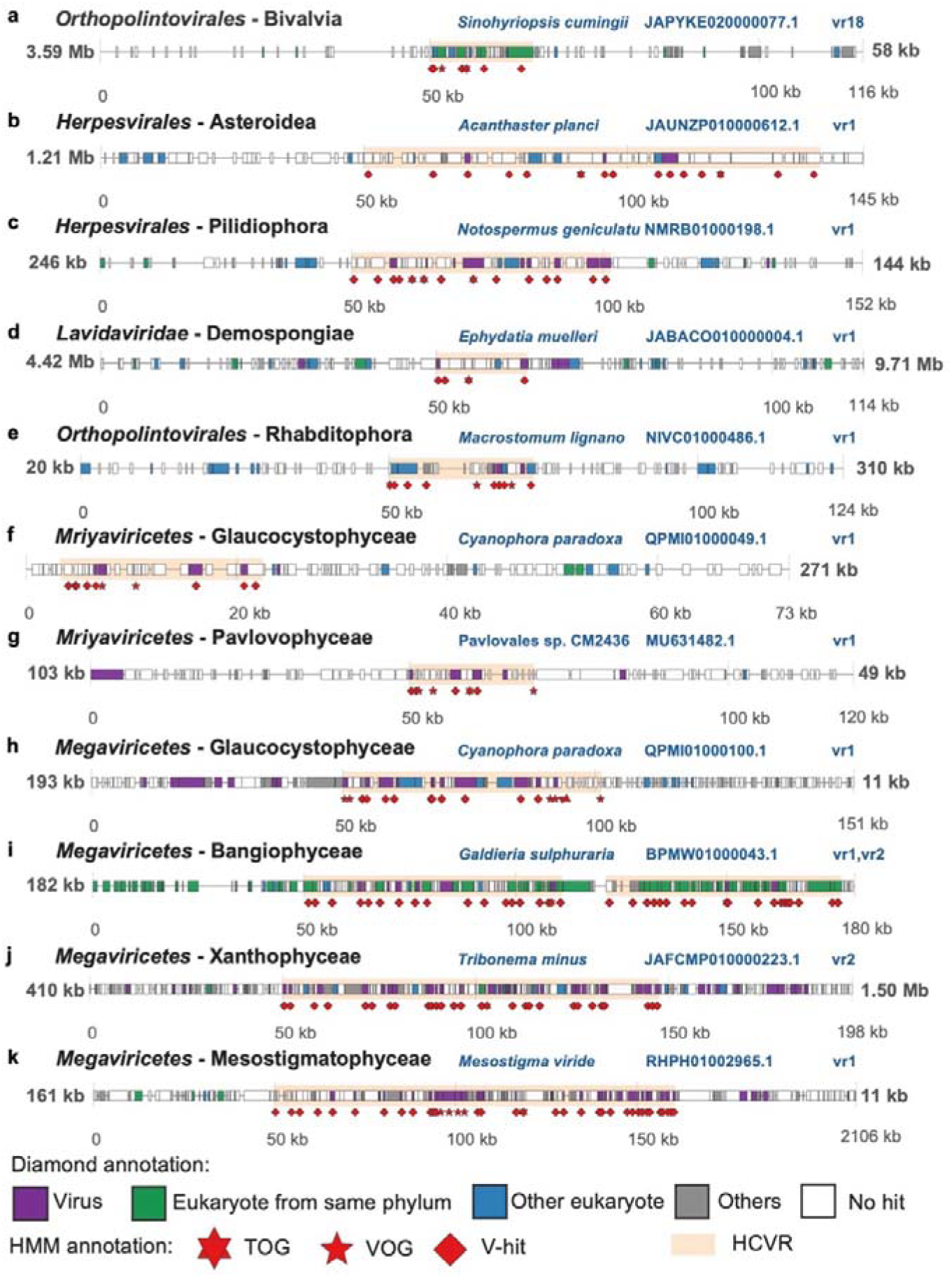
Embedded HCVRs in eukaryotes. For each map, 50 kb of flanking sequence on both sides of each HCVR is shown; when the available flanking sequence was longer than 50 kb, the unshown length is indicated as text. HCVRs are highlighted with a light-red background. Each rectangle represents an ORF. Rectangle colors correspond to the taxonomic affiliations of the best hits. HMM-based annotations are shown below each map.

Current knowledge of BC I viruses in invertebrates remains limited and is strongly biased toward arthropods, molluscs, and cnidarians (**Fig. 4**). For example, herpesviruses are known to infect molluscs and have also been putatively associated with annelids^36^. In our dataset, *Herpesvirales*-related HCVRs were embedded within the genomes of a starfish and a ribbon worm (**Fig. 5b, c**). These HCVRs encoded DNA polymerases, and phylogenetic analysis placed these sequences in lineages branching between the *Orthoherpesviridae* and *Malacoherpesviridae* clades (**Supplementary Fig. S2**), suggesting the existence of divergent herpesviral lineages. We further identified a *Lavidaviridae*-related HCVR embedded in a sponge chromosome (**Fig. 5d**), whereas lavidavirids are currently known to infect protists in association with their helper imiterviruses (*Megaviricetes*). In addition, *Orthopolintovirales*-related insertions were detected in Rhabditophora (flatworm), with most ORFs showing best hits to proteins annotated as other eukaryotes or viruses (**Fig. 5e**). Because closely related eukaryotic genomes of these lineages are poorly represented in current reference databases, sequence-similarity-based annotations alone may underestimate or ambiguously classify such elements more frequently than in bivalves (**Fig. 5a**). The embedded genomic context of these HCVRs therefore provides important evidence for previously overlooked virus–host associations. Together, these examples suggest that the virosphere of invertebrates remains substantially underexplored.

Beyond metazoans, embedded HCVRs also revealed diverse virus-related signals in algal lineages from distinct supergroups. We identified *Mriyaviricetes*-related HCVRs in a Glaucocystophyceae assembly (**Fig. 5f, Supplementary Fig. S2**). *Mriyaviricetes* represents a recently recognized viral clade related to *Pleurochrysis* sp. endemic viruses, which infect members of Prymnesiophyceae, a distinct class of Haptophyta. We further detected the detection of a *Mriyaviricetes*-related HCVR in a Pavlovophyceae genome (**Fig. 5g**). *Megaviricetes*-related HCVRs were found to be embedded in diverse algal genomes, including Bangiophyceae (**Fig. 5h**), Xanthophyceae (**Fig. 5i, Supplementary Fig. S2**), and Mesostigmatophyceae (**Fig. 5j**).

Although endogenization may occur incidentally and does not necessarily imply an active contemporary host–virus relationship^2,45^, the diversity and relative completeness of HCVRs analyzed here (**Figs. 4 and 5**) suggest that these interactions may not date back to the distant past.

### Lineage-specific patterns suggest distinct routes of viral genome endogenization

Phylogenetic analyses combining reference viral sequences and HCVR-derived sequences showed that many HCVRs occupied branches distinct from those of currently sampled viruses (**Supplementary Fig. S2**). The distribution of integration-like VRs was highly uneven across viral major taxa. Very few such signals were detected for *Adenoviridae* and *Papovaviricetes*, whereas integration-like HCVRs assigned to *Lavidaviridae*, *Megaviricetes*, *Mriyaviricetes*, and PLVs were more abundant. An imbalanced distribution of integration-like VRs was also evident within several viral taxa. For example, HCVRs were concentrated in deep branching lineages of *Herpesvirales*. HCVRs from *Mirusviricota* were not identified in the clade of viruses detected in marine ecosystems (i.e., the *Demutovirales*^46^), but were mainly found in other orders. *Orthopolintovirales* VRs were widely distributed across the tree, except for one clade composed exclusively of reference sequences derived from aquatic environmental metagenomes. Together, these patterns suggest a partial separation between the contemporary observable virosphere and the endo-virosphere inferred from eukaryotic genomic data.

To gain insight into the mechanism of viral insertion, we examined genes potentially related to mobility or genome integration (**Supplementary Fig. S4**). In PLV-related and *Orthopolintovirales*-related elements, mobility-related genes appeared more frequently in HCVRs than in isolated or metagenome-derived reference viral genomes (**Supplementary Fig. S5a, b**), and a similar tendency was also observed for *Herpesvirales* (**Supplementary Fig. S5c**). These patterns suggest that viral lineages differ in their propensity to leave traces in eukaryotic genomes, even within the same viral major taxon. For PLVs and *Orthopolintovirales*, lineage-specific mobility modules may contribute to the abundance of HCVRs. For *Herpesvirales*, chromosomal integration mediated by telomeric repeats is known in specific lineages, such as human herpesvirus 6 and avian alphaherpesviruses^9,47^. The detection of mobility-related genes in HCVRs from deep-branching *Herpesvirales* lineages suggests that these lineages may represent integration-prone herpesviruses. By contrast, in the three major viral taxa within *Nucleocytoviricota* (**Table 1**), mobility-related genes were encoded by some reference viral genomes but were less frequently detected in HCVRs (**Supplementary Fig. S5d–f**). *Nucleocytoviricota*-related HCVRs lacking mobility-related genes may therefore mainly represent accidental insertions associated with active viral replication, possibly mediated by virally encoded mobility genes that were not retained within the detected HCVRs.

Differences in replication compartments, infection strategies, host range, and opportunities for interaction with mobile genetic elements could all influence whether a viral lineage becomes detectable as EVEs. Viral life cycles involving chromosomal integration have emerged sporadically across viral evolution, as exemplified by phaeoviruses such as *Ectocarpus siliculosus* virus and the recently characterized integrated giant virus in a green alga^11,13^. Even within the same viral group, the subcellular site of replication can differ among lineages, as suggested for mirusviruses^46^. The imbalanced phylogenetic distributions of VRs across viral trees may therefore reflect lineage-specific differences in viral life cycle, replication compartments, and opportunities for interaction with mobile genetic elements. Together, these observations suggest that some dsDNA viral lineages may be more prone than others to leave detectable traces in eukaryotic genomes.

## Conclusion

Systematic mining identified BC I VRs in a broad spectrum of eukaryotic lineages, with particularly pronounced signals in invertebrates. As sequence-based surveys increasingly reveal viral diversity prior to isolation or microscopic observation of viruses, the putative associations and embedded viral signals presented in this study provide entry points for exploring viruses across poorly sampled eukaryotic lineages. Together, the exceptionally high VR content observed in invertebrates and the embedded HCVRs reported here suggest that invertebrates are reservoirs of hidden dsDNA viral diversity.

The present study is inevitably constrained by the availability of both sequenced eukaryotic genomes and reference viral genomes. In particular, the limited taxonomic coverage restricts the resolution of viral taxonomic assignments. As additional viral groups are discovered and characterized, more genomic regions may be recognized as virus-derived, and some existing assignments may be refined. Resolving virus–host relationships at finer taxonomic scales will require further analytical efforts. Although these issues represent limitations of the present study, the VR catalog generated here provides a foundation for future studies of the endogenous dsDNA virosphere and its role in the evolution of eukaryotic genomes.

## Methods

### Construction of cellular reference HMMs

Cellular reference HMMs were constructed in two sequential steps. First, protein sequences covering 778 eukaryotic species from different genera (one species per genus) were downloaded from the KEGG database^48^. Sequences shorter than 50 amino acids or longer than 4,000 amino acids were removed. OGs were identified using OrthoFinder v2.5.5^49^. OGs present in at least 60% of genomes within at least one of the four major eukaryotic groups (animals, green plants, fungi, or protists) were retained, yielding 12,555 OGs. For each OG, up to 200 sequences were randomly selected and aligned using MUSCLE v5.1^50^. Alignments were trimmed with trimAl v1.4.1^51^ using the “-gt 0.05” option, and trimmed alignments were used to construct HMMs with HMMER v3.4^52^.

The second step was based on HMMs from Pfam-A v37 database^53^. To minimize potential viral contamination, HMMs containing virus-related keywords (i.e., ‘virus’, ‘viral’, ‘phage’, ‘capsid’, ‘double jelly roll’, ‘terminase’) were excluded. The remaining HMMs were then evaluated using hmmscan (E-value < 10 ³) against both the collected viral protein set and eukaryotic proteins from KEGG. HMMs that produced more hits to viral sequences than to eukaryotic sequences were removed, leaving 19,991 HMMs. In total, 32,546 HMMs from both steps were retained as cellular reference models.

### Construction of VOGs and TOGs

We collected reference genomes of the 11 eukaryote-infecting BC I viral major taxa from various sources. For six viral taxa (*Adenoviridae*, *Herpesvirales*, *Lavidaviridae*, *Megaviricetes*, *Naldaviricetes*, and *Papovaviricetes*), we obtained protein sequences from RefSeq genomes in NCBI Virus (as of June 10, 2024). For *Mirusviricota*, *Mriyaviricetes*, PLVs, *Pokkesviricetes*, and *Orthopolintovirales*, we additionally gathered genomic data from other sources (see details in Supplementary Table S2). Gene prediction was performed for large/giant viruses using Prodigal v2.6.3^54^ with default options and for smaller viruses using GeneMarkS v4.30^55^ with the metagenomic model and other default options. For each viral taxon, we identified OGs using Proteinortho v6.3.2^56^ with parameters following a previously described approach^57^ (-e=1e-3 -identity=15 -p=blastp+ -selfblast -cov=40). For each taxon, we constructed viral HMMs based on OGs shared by at least three genomes using the methodology as described in the previous section.

To identify OGs exhibiting viral specificity (i.e., VOGs), we performed hmmsearch using both cellular reference HMMs and viral HMMs against proteins from all OGs identified in viral genomes. An OG was classified as a VOG if at least one protein within the OG had an E-value smaller than 10 against viral HMMs. For all proteins in this OG, the minimum E-value for the cellular reference HMMs had to exceed 10 and be at least 10³ times greater than that of the corresponding viral HMM, ensuring viral specificity. The classification of TOGs was based on VOGs. Proteins within a TOG were required to exhibit the best HMM match to viral HMMs from the corresponding taxon, whereas the minimum E-values for HMMs from other viral major taxa had to exceed 10 and be at least 10³ times higher than those of their own taxon, thereby confirming taxonomic specificity.

### Detection of associations between viral major taxa and eukaryotic contigs

We first retrieved eukaryotic taxonomic information using the Bio.Entrez module of Biopython^58^, followed by manual curation. For each eukaryotic assembly, Prodigal (with configurations adjusted for long input sequences) and GeneMarkS were used to identify ORFs. The existence of introns was not considered here, as the goal of this study was to identify viral-like regions. Thus, the ORF density of a eukaryotic assembly did not reflect the true coding density. We then performed hmmsearch to identify ORFs with significant similarity to VOGs. To inclusively capture candidate contigs in the initial screening step, an E-value threshold of 10 ^3^ was applied. Because some ORFs may match multiple VOGs, only the best hit was considered in each case.

We conducted a secondary hmmsearch using the -Z parameter to standardize E-value calculations across searches, with database sizes specified for each eukaryotic group (Metazoa: 4,962,163; Fungi: 193,281; Viridiplantae: 237,367; Protists: 317,655). If an ORF had a hit to a cellular model with an E-value less than 100 times that of its best VOG match, the hit was removed as a potential cellular sequence. The remaining ORFs with VOG hits were considered viral-like ORFs. A viral taxon–contig pair was recorded if the contig contained either (i) at least one viral-like ORF with a TOG hit or (ii) two viral-like ORFs with hits to distinct VOGs from the viral major taxon. If a virus–contig pair was detected based on the ORFs predicted by either Prodigal or GeneMarkS, the corresponding ORF set was used for downstream analyses. When the same virus–contig pair was detected based on ORFs predicted by both methods, only one ORF set was retained. The method showing better overall performance was selected, defined as detecting a greater number of contigs specifically associated with the corresponding viral major taxon and eukaryotic kingdom.

### Determination of viral regions

We performed hmmsearch on all ORFs within each contig against HMMs constructed from OGs of the corresponding viral taxon and cellular references HMMs (E-value < 10 ³) to obtain viral and cellular annotations. VRs were identified on the basis of these annotations and the genomic locations of ORFs. A region was classified as a VR if it contained multiple distinct viral hits originating from a single viral taxon. Specifically, at least two distinct viral hits were required for *Adenoviridae*, *Lavidaviridae*, *Mriyaviricetes*, PLVs, *Papovaviricetes*, and *Orthopolintovirales*, whereas at least three distinct viral hits were required for *Herpesvirales*, *Megaviricetes*, *Mirusviricota*, *Naldaviricetes*, and *Pokkesviricetes*. In both cases, all viral hits had to be assigned to the dominant viral major taxon within the region. Additionally, no more than 10 ORFs with cellular annotations were allowed between two viral ORFs, and the genomic distance between them could not exceed 10 kb. The length of each VR was determined from the start of the first ORF with a viral best hit to the end of the last ORF with a viral best hit. Viral–cellular annotation tables were generated for each VR based comparing the minimum E-values of hits to cellular reference HMMs and viral HMMs from OGs within the corresponding viral taxon.

In this study, VRs are operationally defined by our detection framework; given the large number of identified VRs, we do not aim to delineate the exact biological units or boundaries of individual VRs, which may be affected by complex evolutionary processes such as rearrangements or recurrent insertions. Because many major viral taxa share homologous genes, preliminary taxonomic assignments based on HMM profiles may not accurately reflect evolutionary relationships.

### Redundancy analysis of VRs in individual assemblies

We extracted nucleotide sequences of VRs from each assembly and clustered them using MMseqs2 v16-747c6^59^ under different thresholds for minimum sequence identity and alignment coverage. Nearly identical VRs were defined using sequence identity and coverage thresholds both set at 0.9.

### TNF composition analysis

For each assembly containing identified VRs, the assembly was divided into viral and cellular components. If the total length of VRs exceeded half the length of the contig on which they were located, the contig was classified as viral. In other cases, the genomic regions adjacent to the VRs within the same contig were treated as cellular components. These regions were combined with contigs lacking VRs to calculate the average cellular TNF vector for each assembly. From the cellular regions, 1,000 random fragments of 3,000 bp were sampled, and their Pearson correlation coefficients with the average cellular TNF vector were calculated to establish an assembly-specific baseline. Pearson correlation coefficients were then calculated between the TNF vector of each VR and the average cellular TNF vector to assess whether the nucleotide composition of VRs differed from that of the cellular components, as described in the main text and Supplementary Fig. S1f.

### Classification of viral regions

Viral contig-like VRs were defined based on two criteria. First, their total length was required to exceed half the length of the corresponding contig. Second, the proportion of annotated ORFs assigned to cellular OGs was required to be less than 20%. VRs that did not simultaneously satisfy both conditions were considered to be potentially integrated into host genomes and were thus classified as integration-like VRs.

To classify HCVRs, we first identified conserved OGs for each viral taxon. Redundant reference viral genomes were removed using vclust v1.3.1 (--metric ani --ani 0.95)^60^. An OG was considered conserved if it was present in more than half of the filtered reference genomes of the corresponding taxon. We then calculated the acceptable range for the proportion of conserved OGs based on the interquartile range (IQR; Q1 − 1.5×IQR to Q3 + 1.5×IQR). An HCVR was required to contain at least one conserved OG, and its proportion of conserved OGs had to fall within the calculated range.

We also evaluated the completeness of each assembly using BUSCO v5.1.2^61^, applying metazoa_odb10, fungi_odb10, viridiplantae_odb10, or eukaryota_odb10 datasets to metazoan, fungal, green plant, and protist assemblies, respectively. VRs from assemblies with completeness scores below 30% were excluded from HCVR classification. Additionally, we calculated the rate of unique hits for each VR using our annotation tables; VRs in which fewer than 50% of hits were unique were excluded.

An HCVR was required to meet two additional conditions: (i) at least 20% of annotated ORFs within the region had their best hits to models constructed from viral sequences, with ≥ 90% of these hits assigned to the same viral major taxon; and (ii) the region contained at least one TOG from its assigned viral major taxon and no TOGs from other taxa.

We performed DIAMOND blastp (v2.1.9, E-value < 10 ^3^)^62^ to compare all ORFs from the HCVR candidates against a custom viral reference database and recorded the dominant taxon of the best hits (**Supplementary Fig. S6**). The custom database was built from the reference viral genomes used in this study and complemented with high-confidence sequences from the corresponding taxa in the IMG/VR v7.1 database^63^. Notably, when this version of the IMG/VR database was compiled, several viral taxa (e.g., *Mirusviricota*, *Mriyaviricetes*, and PLVs) had not yet been formally established. Therefore, TOGs from these taxa were used to identify potentially related misclassified sequences in the database. Specifically, some sequences likely belonging to *Mirusviricota* or *Mriyaviricetes* were annotated as *Nucleocytoviricota* in the database, whereas PLV-related sequences were annotated as *Polisuviricotina*. These sequences were removed from the database prior to analysis.

### Distributional statistics of HCVRs in distinct eukaryotic groups

The information on virus–host associations in specific eukaryotic lineages was obtained from the Virus–Host database (https://www.genome.jp/virushostdb). All associations supported solely by viral contigs were manually examined. As a result, the *Papovaviricetes*–Magnoliopsida association was removed, as the only supporting evidence was a short viral contig (CBTL0109906317.1) showing clear similarity to human papillomavirus, whereas the corresponding host assembly (GCA_900067735.1) was annotated as “contaminated” in GenBank.

### DIAMOND annotation for genomic maps

ORFs encoded on HCVR-containing contigs were searched using DIAMOND with an E-value threshold of 0.1. Each ORF set was queried against two protein databases: the RefSeq protein database^64^ and the previously described curated viral protein database used for taxonomic assessment. For each ORF, the best hit across the two databases was selected based on the highest bit score.

### Phylogenetic analysis of discovered HCVRs and reference viruses

For trees of each major viral taxon, seed sequences were selected as follows, with different functional marker genes chosen depending on the lineage: *Adenoviridae*: YP_002213842.1; *Herpesvirales*: YP_010084960.1; *Lavidaviridae*: YP_002122381.1; *Megaviricetes*: YP_003986825.1; *Mirusviricota*: TARA_AON_NCLDV_00032_000000000015; *Mriyaviricetes*: Yaravirus_MCP_consensus; *Naldaviricetes*: YP_009116659.1; *Papovaviricetes*: NP_041327.2; PLVs: SAF1_17; *Pokkesviricetes*: YP_009162448.1; *Orthopolintovirales*: Polinton-1_DY. JackHMMER^65^ was used to identify homologs of these marker genes. For each lineage, reference proteins were pooled into a lineage-specific database, which was searched using the corresponding seed sequence for five iterations. Redundant sequences were removed using CD-HIT v4.8.1^66^, with lineage-specific thresholds for *Orthopolintovirales* (-c 0.6 -aS 0.6 -n 4) and PLVs (-c 0.8 -aS 0.8 -n 5). These thresholds were chosen to retain sufficient representation of homologous sequences while reducing redundancy for downstream phylogenetic analyses.

The dereplicated HCVR-derived sequences were then combined with reference sequences. Multiple sequence alignments were performed using Clustal Omega v1.2.4^67^ and trimmed with trimAl using a gap threshold of 0.05. Aligned sequences in which gaps accounted for more than 90% of sites were removed. Phylogenetic trees were reconstructed using IQ-TREE v2.2.0^68^ with 1,000 ultrafast bootstrap replicates^69^. Models were selected with ModelFinder^70^. The concatenated phylogenetic trees for *Megaviricetes* and *Orthopolintovirales* were reconstructed using the same procedure, except that gap-rich sites were not removed. The phylogenetic trees were visualized with iTOL v7^71^.

### Statistics of mobility-related genes

To assess the distribution of genes potentially related to genome mobility or integration, ORFs encoded within VRs and reference viruses were screened using functional annotations from eggNOG-mapper v2.1.12^72^. ORFs annotated as integrase / *rve*, recombinase, transposase, resolvase, or endonuclease were identified by case-insensitive keyword matching of annotation fields. The presence or absence of each gene category was then summarized for each VR or reference viral genome.

## Acknowledgements

This work was supported by JSPS/KAKENHI (nos. 25KJ1513 and 22H00384), iJURC (2024-27) and HFSP (Ref. No.: RGP011/2024). Computational time was provided by the Supercomputer System, Institute for Chemical Research, Kyoto University (Kyoto, Japan). The authors thank Edanz (https://jp.edanz.com/ac) for editing a draft of this manuscript. The authors thank Dr. Kenji K. Kojima from Genetic Information Research Institute for his valuable suggestions.

## Author Contributions

HZ designed the study, performed most of the analyses and wrote the first draft of the manuscript. LM contributed to the phylogenetic analyses. RZ contributed to the gene function analyses. MG contributed to the investigation of individual VRs. HO supervised the study and contributed to the design of the bioinformatics pipeline. All authors have read and approved the final version of the manuscript.

## Data Availability

Table for VRs, HMM files of VOG/TOG and their annotations, sequences and annotation files of VRs, multiple sequence alignments for phylogenetic analysis, tree files, and annotation files for visualization in iTOL (https://itol.embl.de/shared/1VKuwR8IMDABB) are available for review via Zenodo.

## Declaration of Interests

The authors declare no competing interests.

## Supplemental dataset legends

**Supplementary Table S1: Eukaryotic assemblies used in this study**

**Supplementary Table S2: Reference viral genomes used in this study**

**Supplementary Table S3: Summary statistics of VRs across major viral taxa**

**Supplementary Table S4: Proportion of redundant VRs across individual assemblies**

**Supplementary Table S5: Previously reported endogenization between eukaryotes and dsDNA viruses**

**Supplementary Data S1: Information on identified VRs**

---

**Supplementary Fig. S1.**
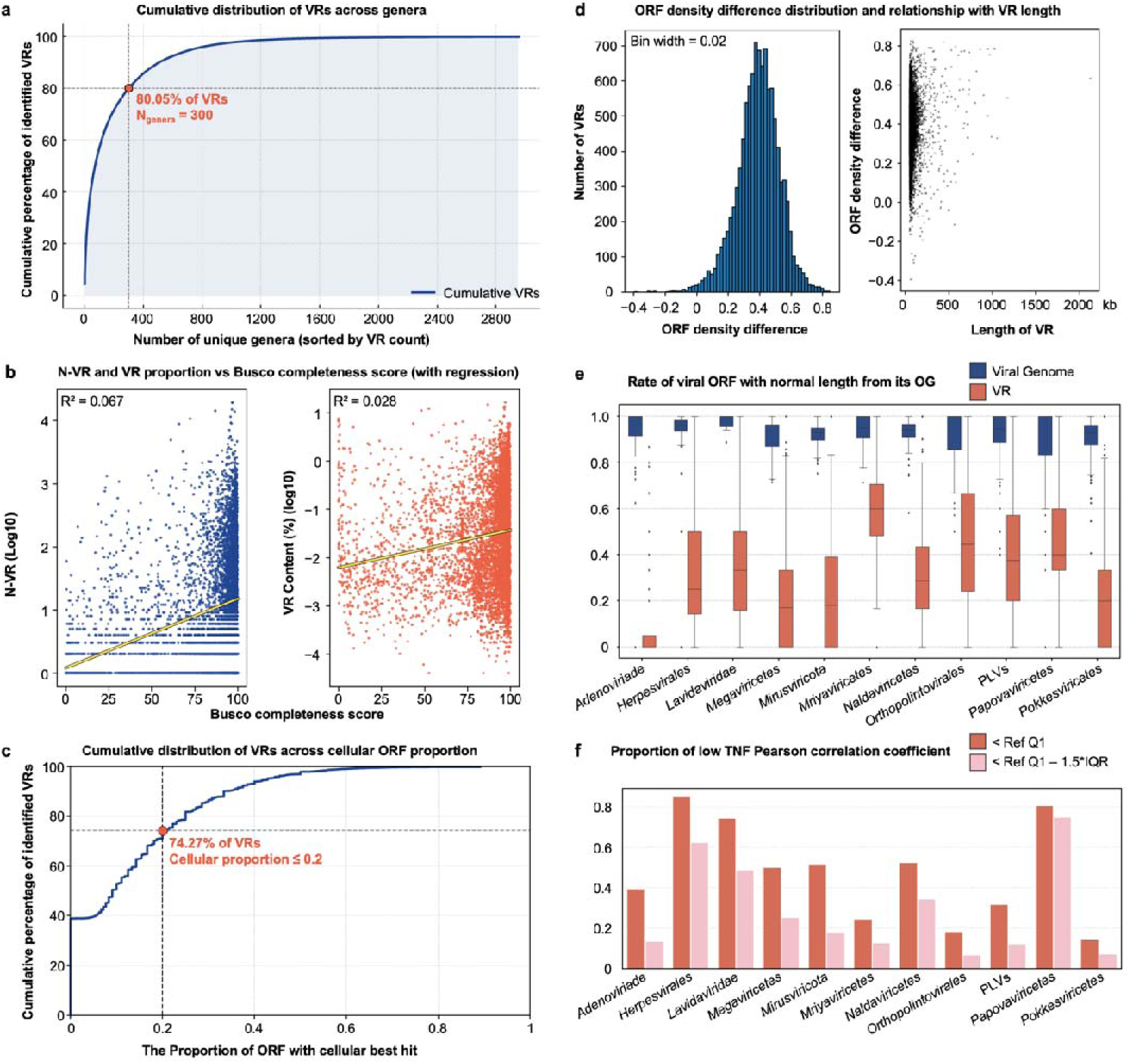
Properties of VRs (a–f) Subfigure titles are shown at the top of each panel. **(b)** The left plot represents the number of VRs, and the right plot represents the proportion of VRs in each assembly. Each dot corresponds to one assembly, and the regression line is shown in yellow. **(d)** The difference represents the ORF density of VRs minus that of the corresponding. Considering the effect of the definition of a VR (i.e., it starts with an ORF and ends with another ORF) on the computed ORF density, we only considered VRs longer than 50 kb. **(e)** The y-axis represents, for each VR or viral genome, the proportion of ORFs within the normal length range of their corresponding OG. **(f)** The y-axis represents, for all VRs of a given taxon, the proportion of VRs whose TNF composition differs from that of the host assembly.

**Supplementary Fig. S2.**
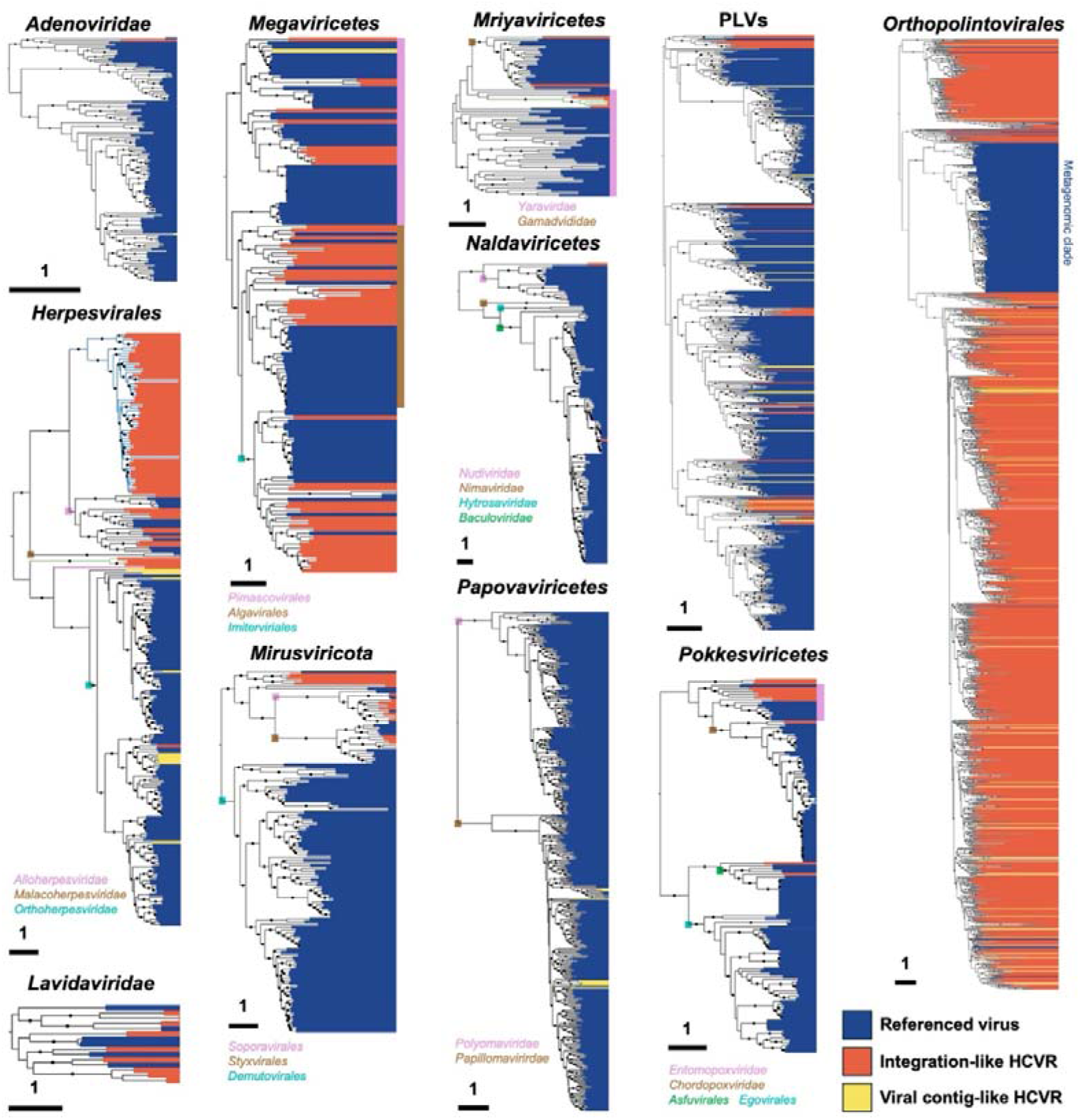
Reconstructed phylogenetic trees of HCVR-derived and viral sequences. Each tree corresponds to a specific viral taxon, as indicated above. Scale bars are shown in the lower left corners, and all trees are midpoint rooted. Black circles indicate bootstrap support values higher than 90. Blue, orange, and yellow shading indicate reference viral sequences, integration-like HCVRs, and viral contig-like HCVRs, respectively. For *Adenoviridae*, *Herpesvirales*, *Megaviricetes*, *Naldaviricetes*, and *Pokkesviricetes*, DNA polymerase sequences were used for tree reconstruction. For *Lavidaviridae*, *Mirusviricota*, *Mriyaviricetes*, and *Orthopolintovirales*, the major capsid protein sequences were used. For *Papovaviricetes* and PLVs, E1 helicase and pATPase were used, respectively. The tree for *Megaviricetes* were manually curated. In the *Herpesvirales* tree, branches associated with Actinopterygii, Asteroidea, and Pilidiophora were colored blue, pink, and green, respectively. In the *Megaviricetes* tree, the Xanthophyceae-associated branch was colored green. In the *Mriyaviricetes* tree, branches associated with Pavlovophyceae and Glaucocystophyceae were colored blue and green, respectively. The selected models for each viral lineage were as follows: *Adenoviridae*: Q.pfam+F+I+G4; *Herpesvirales*: Q.pfam+F+I+G4; *Lavidaviridae*: Q.pfam+F+G4; *Megaviricetes*: VT+F+G4; *Mirusviricota*: Q.pfam+F+I+G4; *Mriyaviricetes*: Q.pfam+F+G4; *Naldaviricetes*: Q.yeast+F+I+G4; *Papovaviricetes*: Q.yeast+F+I+G4; PLVs: Q.pfam+F+I+G4; *Pokkesviricetes*: VT+F+I+G4; and *Orthopolintovirales*: VT+F+G4.

**Supplementary Fig. S3.**
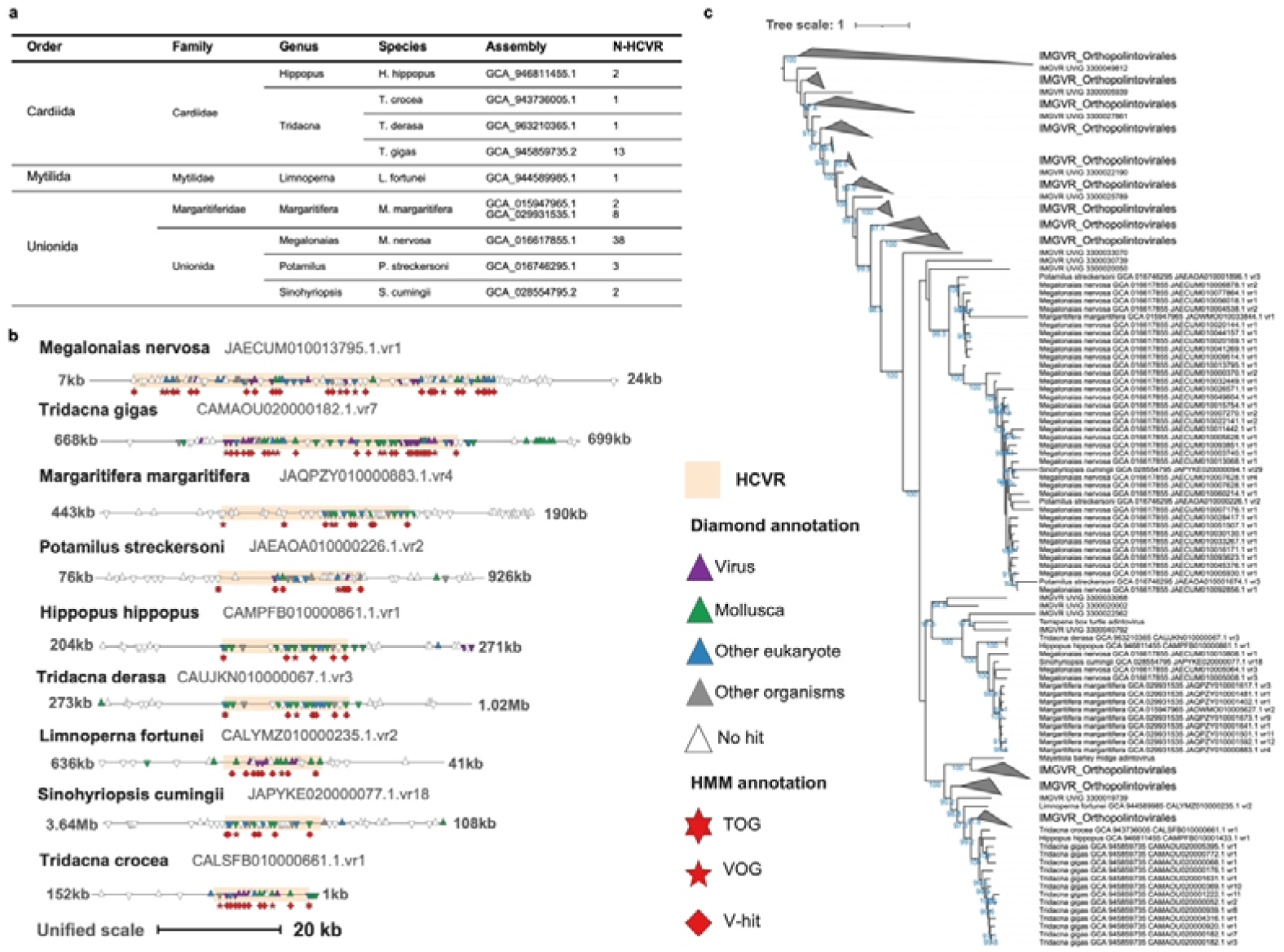
HCVRs assigned to *Orthopolintovirales* in bivalve assemblies. **(a) Taxonomic information and numbers of *Orthopolintovirales*-related HCVRs in bivalve assemblies. (b) Genomic maps of HCVRs in each bivalve assembly.** For each map, 20 kb of flanking sequence on both sides of each HCVR is shown; when the available flanking sequence is longer than 20 kb, the unshown length is indicated as text. HCVRs are highlighted with a light-red background. Each triangle represents an ORF; upward and downward triangles indicate ORFs encoded on the positive and negative strands, respectively. Triangle colors correspond to the taxonomic affiliation of the best hit. HMM-based annotations are shown below each map. **(c) Reconstructed phylogenetic tree of HCVR-derived and reference *Orthopolintovirales* sequences.** The tree was inferred from a concatenated alignment of five OGs: Orthopolintovirales_og1, og2, og5, og23, and og33. The best-fit model was VT+F+I+G4.

**Supplementary Fig. S4.**
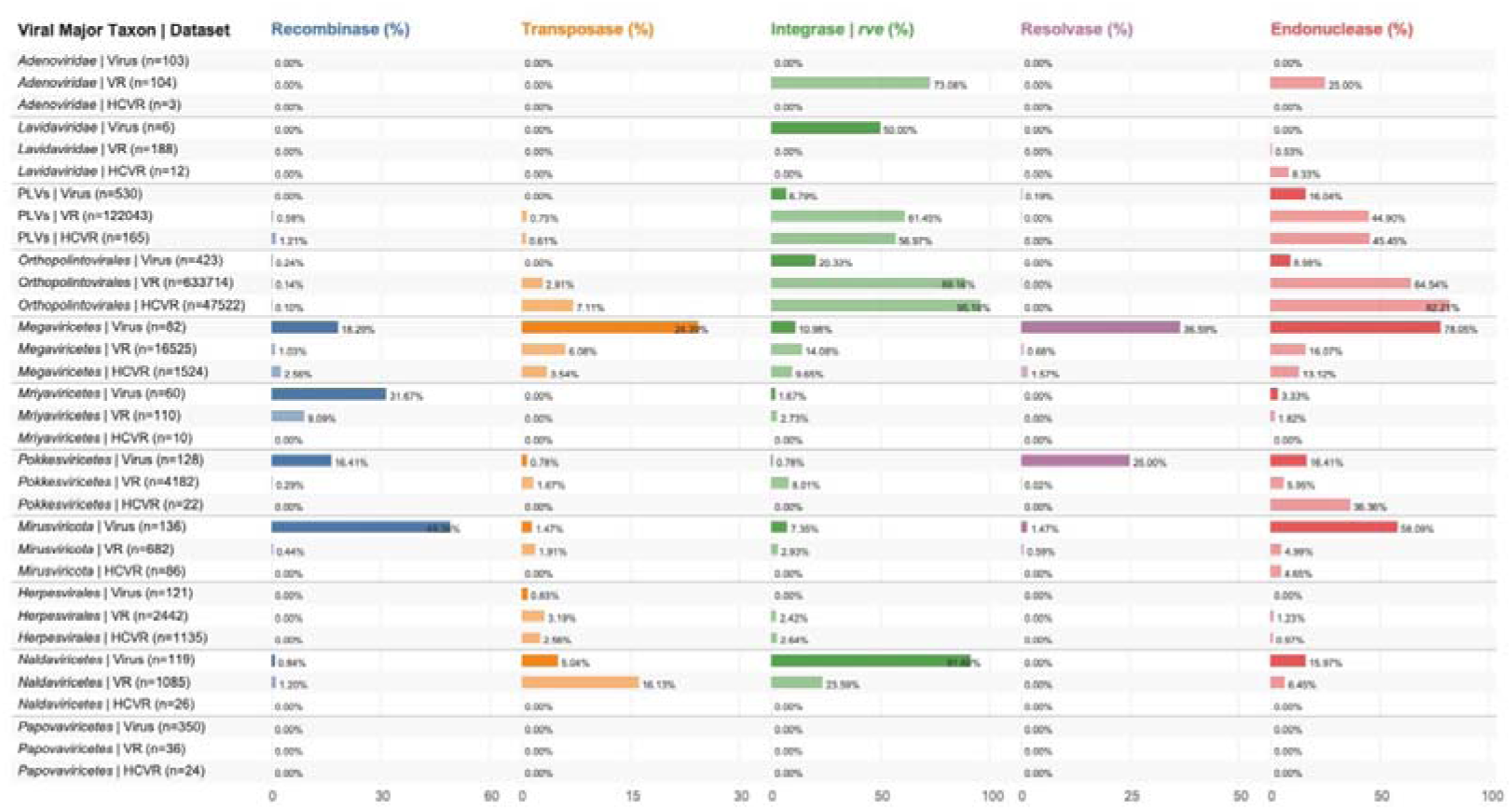
Distribution of mobility-related genes across reference viruses, VRs, and HCVRs. For each viral major taxon, the proportions of entries encoding genes potentially related to mobility or genome integration are shown for three datasets: reference viruses, VRs, and HCVRs. Gene categories include recombinase, transposase, integrase/*rve*, resolvase, and endonuclease. Numbers in parentheses indicate the number of entries analyzed for each viral major taxon and dataset. Bars indicate the percentage of entries containing at least one ORF assigned to each gene category.

**Supplementary Fig. S5.**
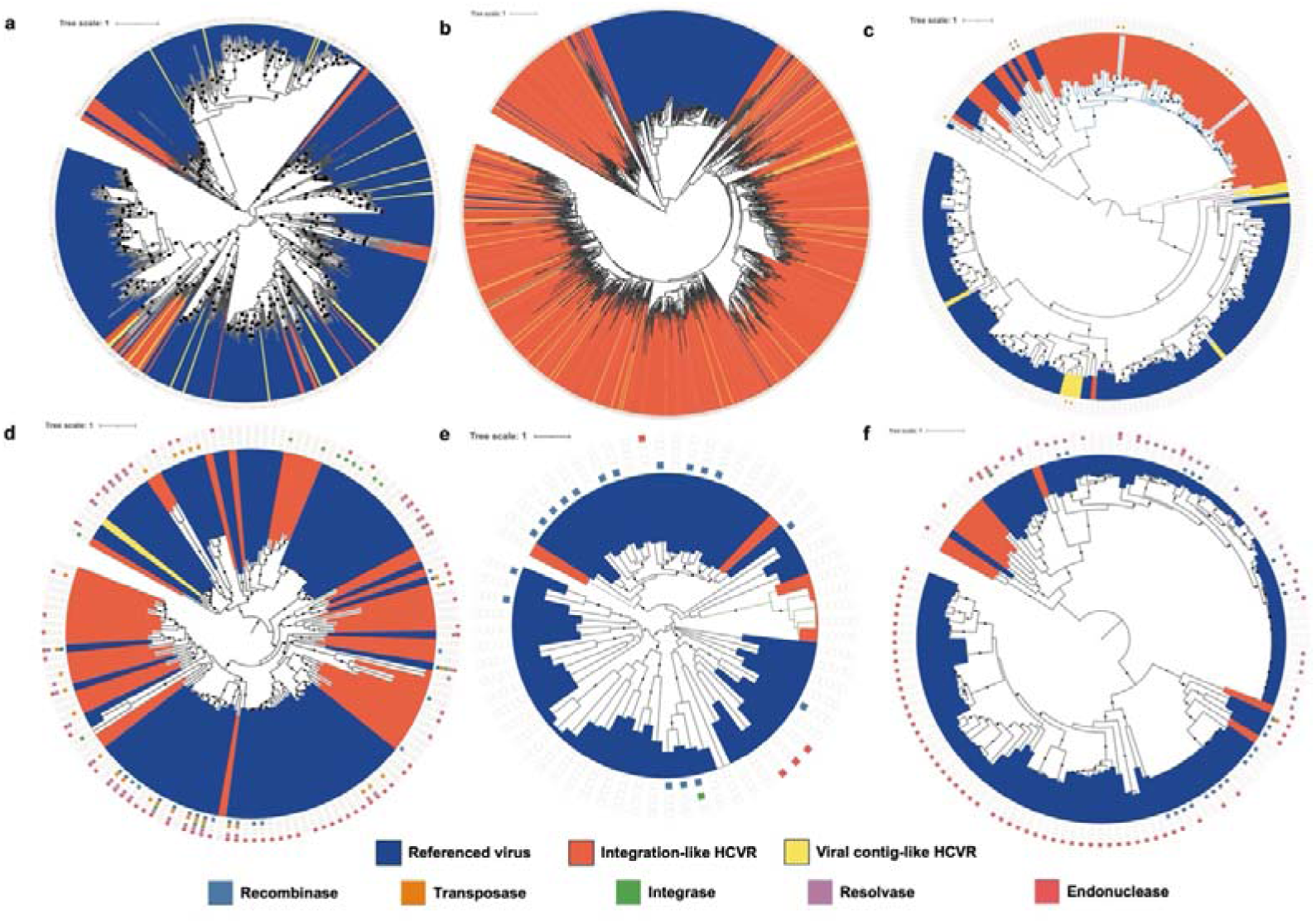
Phylogenetic distribution of mobility-related genes across selected viral major taxa. Phylogenetic trees are shown for selected viral major taxa, corresponding to those in Supplementary Fig. S2: **(a)** PLVs; **(b)** *Orthopolintovirales*; **(c)** *Herpesvirales*; **(d)** *Megaviricetes*; **(e)** *Mriyaviricetes*; and **(f)** *Pokkesviricetes*. Branch-associated color blocks indicate reference viruses, integration-like HCVRs, and viral contig-like HCVRs. Outer symbols indicate the presence of mobility-related gene categories. An interactive version of the trees is available (see Data Availability).

**Supplementary Fig. S6.**
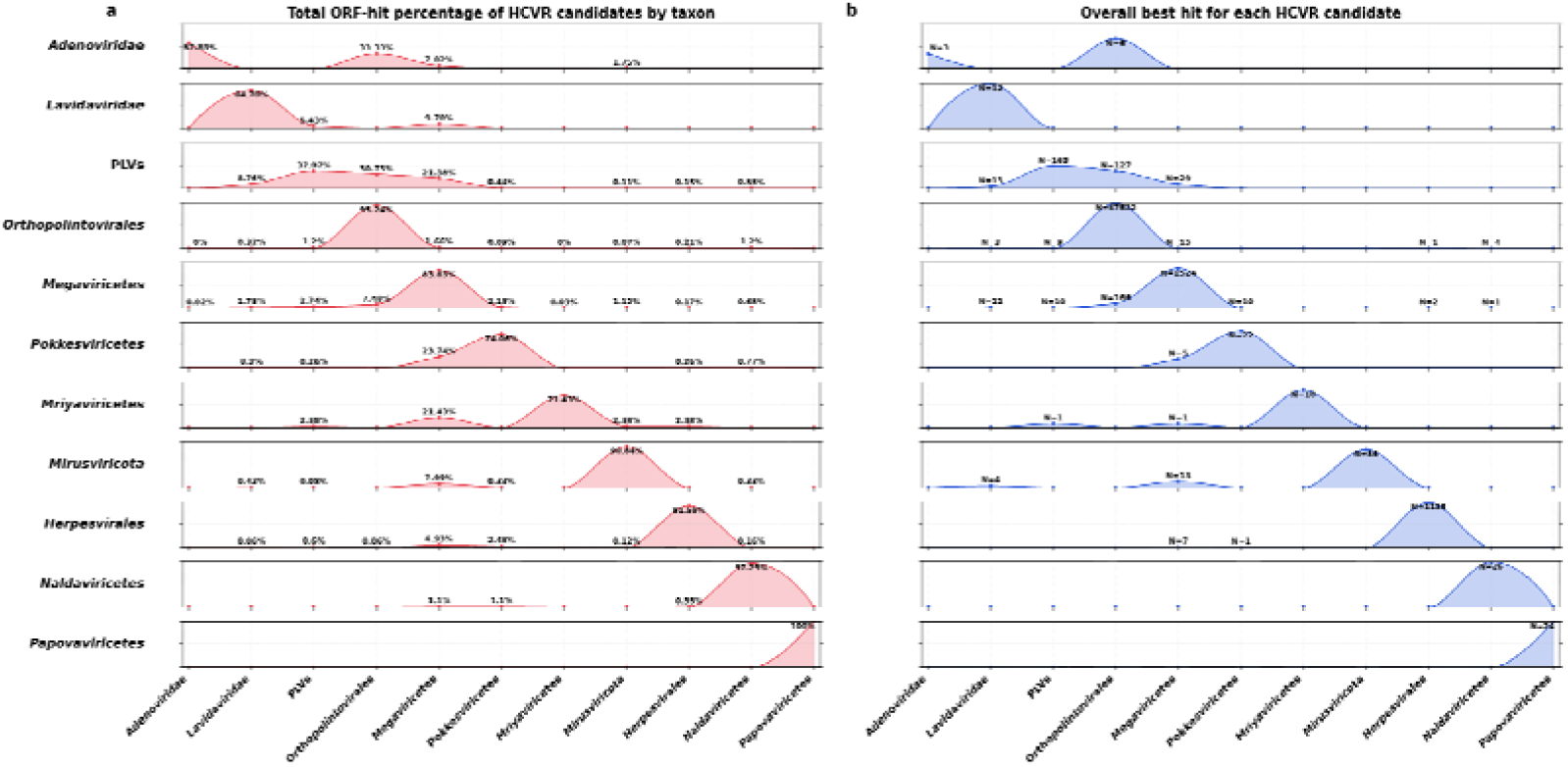
Independent taxonomic assessment of HCVR candidates. Each row corresponds to filtered candidates assigned to one viral major taxon in our marker-based classification. **(a) Taxonomic distribution of all gene-level hits detected in HCVR candidates belonging to each viral taxon.** The vertical axis indicates the proportion of hits assigned to each reference viral taxon, and the horizontal axis represents viral taxa in the expanded reference database. **(b) Taxonomic assessment based on the overall best-hit taxon for each HCVR candidate.** For each HCVR, hits from all encoded genes were summarized, and the viral taxon receiving the highest cumulative support was assigned as the overall best-hit taxon. The plots show the proportion of HCVRs whose overall best-hit taxon corresponded to each viral taxon in the reference database.

## Notes

### Competing Interest Statement

The authors have declared no competing interest.

### Summary of Updates

This revised version substantially updates the manuscript structure, title, abstract, figures, and interpretation. We reframed the study as a research article focused on lineage specific accumulation of endogenous dsDNA viral elements across eukaryotes. The revised manuscript includes updated analyses of viral region distribution across host lineages and viral taxa, a more stringent definition of taxonomically high confidence viral regions, and a revised assessment of putative virus host associations. We also added and revised genomic examples supporting endogenization, including cases involving invertebrates, diverse algae, and Megaviricetes related viral regions in ray finned fishes. The figures and supplementary materials have been reorganized to better support the revised narrative, and the text has been edited throughout for clarity, consistency, and more conservative interpretation.

